# Metagenome-genome-wide association studies reveal human genetic impact on the oral microbiome

**DOI:** 10.1101/2021.05.06.443017

**Authors:** Xiaomin Liu, Xin Tong, Jie Zhu, Liu Tian, Zhuye Jie, Yuanqiang Zou, Xiaoqian Lin, Hewei Liang, Wenxi Li, Yanmei Ju, Youwen Qin, Leying Zou, Haorong Lu, Xun Xu, Huanming Yang, Jian Wang, Yang Zong, Weibin Liu, Yong Hou, Shida Zhu, Xin Jin, Huijue Jia, Tao Zhang

## Abstract

The oral microbiota contains billions of microbial cells, which could contribute to diseases in a number of body sites. Challenged by eating, drinking and dental hygiene on a daily basis, the oral microbiota is regarded as highly dynamic. Here, we report significant human genomic associations with the oral metagenome from more than 1,915 individuals, for both the tongue dorsum and saliva. We identified five genetic loci associated with oral microbiota at study-wide significance (*p* < 3.16 × 10^−11^). Four of the five associations were well replicated in an independent cohort of 1,439 individuals: rs1196764 at *APPL2* with *Prevotella jejuni, Oribacterium uSGB 3339* and *Solobacterium uSGB 315*; rs3775944 at the serum uric acid transporter *SLC2A9* with *Oribacterium uSGB 1215, Oribacterium* uSGB 489 and *Lachnoanaerobaculum umeaense*; rs4911713 near *OR11H1* with species *F0422 uSGB 392;* and rs36186689 at *LOC105371703* with *Eggerthia*. Further analyses confirmed 84% (386/455 for tongue dorsum) and 85% (391/466 for saliva) of genetic-microbiota associations including 6 genome-wide significant associations mutually validated between the two niches. Human genetics accounted for at least 10% of oral microbiome compositions between individuals. Machine learning models showed that polygenetic risk score dominated over oral microbiome in predicting predisposing risk of dental diseases such as dental calculus and gingival bleeding. These findings indicate that human genetic differences are one explanation for a stable or recurrent oral microbiome in each individual.

## Introduction

A health individual swallows 1-1.5 liters of saliva every day^1^, the microbes in which could colonize the gut of susceptible individuals^2-5^. Oral metagenomic shotgun sequencing data has been available from the Human Microbiome Project (HMP)^6^, for rheumatoid arthritis^7^ and colorectal cancer^3^. Other diseases such as liver cirrhosis, atherosclerotic cardiovascular diseases, type 2 diabetes, and colorectal cancer studied by metagenome-wide association studies (MWAS) using gut microbiome data also indicated potential contribution from the oral microbiome in disease etiology^2,8-11^.

Controversy over human genetic versus environmental determination of the fecal microbiome is being clarified by an increasing number of studies^12-17^. The strongest signal in cohorts of European ancestry is the association between *LCT1* and *Bifidobacterium*, explained by metabolism of lactose by the commensal bacterium. These large-scale genome-wide association studies have mainly focused on fecal microbiome, however, the influence of host genetics on the composition and stability of the oral microbiome is still poorly understood. Several studies based on 16S rRNA amplicon sequencing and microarrays have reported that human oral microbiota is influenced by both host genetics and environmental factors^18-21^. Only two studies have identified limited human genes that affected oral microbial communities. One study identified that *IMMPL2* on chromosome 7 and *INHBA-AS1* on chromosome 12 could influence microbiome phenotypes^19^. The other study reported a gene copy number (CN) of the *AMY1* locus correlated with oral and gut microbiome composition and function^22^. These two studies used 16S rRNA amplicon sequencing for a small number of samples. Therefore, the influence of human genes on the composition of the oral microbiome and genetic stability between different oral niches are still poorly understood.

Here, we presented the first large-scale metagenome-genome-wide association studies (M-GWAS) in a cohort of 2,984 healthy Chinese individuals, of which all individuals had high-depth whole genome sequencing data and over 1,915 individuals had matched tongue dorsum and salivary samples with metagenomic sequencing data. We further validated the identified associations in an independent replication cohort of 1,494 individuals with also metagenomic sequencing data and relatively low-depth whole genome sequencing data. A large number of concordant associations were identified between genetic loci and the tongue dorsum and salivary microbiomes. The effects of environmental factors and host genes on oral microbiome composition were investigated. Host genetics explained more variance of microbiome composition than environmental factors. The findings underscore the value of M-GWAS for *in situ* microbial samples, instead of focusing on feces.

## Results

### The oral microbiome according to metagenomically assembled microbial genomes

The 4D-SZ cohort (multi-omics, with more data to come, from Shenzhen, China) at present have high-depth whole-genome sequencing data from 2,984 individuals (mean depth of 33×, ranging from 15× to 78×, **Supplementary Table 1, Supplementary Fig. 1**). Among these, over 1,915 individuals had matched tongue dorsum and salivary samples for M-GWAS analysis.

Shotgun metagenome sequencing was performed for the 3,932 oral samples, with an average sequencing data of 19.18 ± 7.90 Gb for 2,017 tongue dorsum and 13.64 ± 2.91 Gb for 1,915 salivary samples (**Supplementary Table 1, Supplementary Fig. 1**). The microbiome composition was determined according to alignment to a total of 56,213 metagenome-assembled genomes (MAGs) that have been organized into 3,589 species-level clusters (SGBs) together with existing genomes, of which 40% (1,441/3,589) was specific in this cohort^23^. Both the tongue dorsum and the salivary samples contained the phyla *Bacteroidetes* (relative abundance of 37.2 ± 11.3% for tongue dorsum and 40.1 ± 10.2% for saliva, respectively), *Proteobacteria* (30.1 ± 16.5% and 30.6 ± 13.1%, respectively), *Firmicutes* (20.5 ± 8.2% and 17.7 ± 6.7%, respectively), *Actinobacteriota* (4.3 ± 3.4% and 2.6 ± 2.0%, respectively), *Fusobacteriota* (4.0 ± 1.9% and 3.3 ± 1.4%, respectively), *Patescibacteria* (in Candidate Phyla Radiation, CPR, 2.5 ± 1.6% and 3.1 ± 1.6%, respectively) and *Campylobacterota* (1.1 ± 0.9% and 1.3 ± 0.8%, respectively) (**Supplementary Fig. 2a-b**).These seven phyla cover between 99.7% (tongue dorsum) and 98.7% (saliva) of the whole community, indicating that the two oral sites share a common core microbiota. Consistent with HMP results using 16S rRNA gene amplicon sequencing^24^, the salivary samples presented a higher alpha diversity than tongue dorsum samples (mean Shannon index of 6.476 vs 6.228; Wilcoxon Rank-Sum test *p*□<□2.2□×□10^−16^; **Supplementary Fig. 2c**). The microbiome compositions calculated by beta-diversity based on genus-level Bray– Curtis dissimilarity slightly differed (explained variance R^2^ = 0.055, *p* < 0.001 in permutational multivariate analysis of variance (PERMANOVA) test; **Supplementary Fig. 2d**).

### Host genetic variants strongly associated with the tongue dorsum microbiome

With this so-far the largest cohort of whole genome and whole metagenome data, we first performed M-GWAS on the tongue dorsum microbiome. With the 1,583 independent tongue dorsum microbial taxa (r^2^ < 0.8 from 3177 taxa total using a greedy algorithm, **Methods**), and 10 million human genetic variants (minor allele frequency (MAF) ≥ 0.5%), 455 independent associations involving 340 independent loci (distance < 1MB and r^2^ < 0.2) and 385 independent taxa reached genome-wide significance (*p* < 5 × 10^−8^). With a more conservative *Bonferroni*-corrected study-wide significant *p* value of 3.16 × 10^−11^ (= 5 × 10^−8^ / 1,583), we identified 3 genomic loci, namely *APPL2, SLC2A9 and MGST1*, associated with 5 tongue dorsum microbial features involving 112 SNP-taxon associations (**Fig. 1a**). These associations showed remarkable evidence of polygenicity and pleiotropy (**Fig. 1b**). There was no evidence of excess false positive rate in the GWAS analysis (genomic inflation factors λ_GC_ ranged from 0.981 to 1.023 with median 1.005; **Supplementary Fig. 3a**). All genome-wide significant associations were listed in **Supplementary Table 2**.

**Fig. 1.**
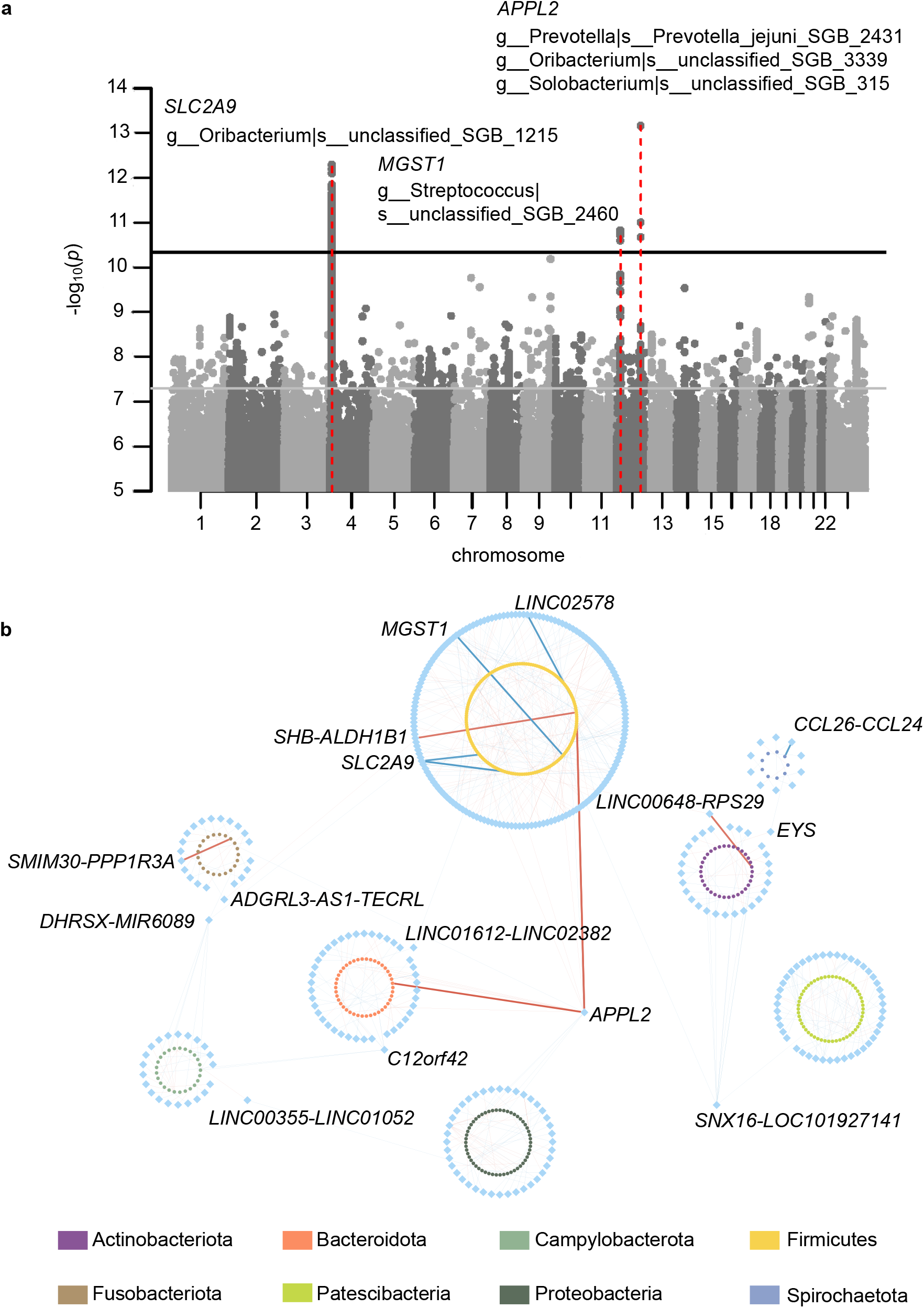
Host genetic signals associated with tongue dorsum microbiome. (**a**). Manhattan plot shows the genetic variants associated with the tongue dorsum microbial taxa. The horizontal grey and black lines represent the genome-wide (*p*□=□5□×□10^−8^) and study-wide (*p*□=□3.16□×□10^−11^ for 1,583 independent M-GWAS tests) significance levels, respectively. Three loci that associated with tongue dorsum microbiome and reached study-wide significance were marked in red. Their located genes and associated microbial taxa with *p* values of < 3.16□×□10^−11^ were also listed. (**b**). Network representation of the 455 gene-microbiome associations identified in the tongue dorsum M-GWAS at the genome-wide significance. Each node represents either a gene (blue diamonds) or a microbial taxon (circles with different colors according to phylum). Each edge is an association between one gene and one microbial taxon. The bold edge represented study-wide significant associations as showed in (**a**). The genes that linked to at least two different microbial taxa from different phyla were also listed.!

We also used a replication cohort to validate these associations. The replication cohort was comprised of 1,494 individuals from multiple cities in China (also shotgun metagenomic sequencing for 1,333 tongue dorsum samples to an average of 19.90 ± 7.73 Gb and 1,299 salivary samples to an average of 13.66 ± 2.80 Gb, but about 9× (ranging from 5× to 15×) whole-genome sequencing for human genome; **Supplementary Table 1, Supplementary Fig. 1**). Among the 455 independent associations identified in the discovery cohort with *p* < 5 × 10^−8^, 33 were not available in the low-depth replication dataset and we were able to replicate 37 associations in the same effect direction of minor allele (*p* < 0.05; **Supplementary Table 2**). Two of the three study-wide signals from the discovery cohort were well replicated: rs1196764 at *APPL2* with *Prevotella jejuni, Oribacterium uSGB 3339* and *Solobacterium uSGB 315*; and rs3775944 at *SLC2A9* with *Oribacterium uSGB 1215, Oribacterium uSGB 489* and *Lachnoanaerobaculum umeaense*.

The strongest association identified by tongue dorsum M-GWAS was on rs1196764 located in the *APPL2* locus, with positive associations with three species, namely *Prevotella jejuni* (**Fig. 2a;** *p*_*discovery*_ = 6.89 × 10^−14^; *p*_*replication*_ =1.00 × 10^−44^), *Oribacterium unclassified SGB (uSGB) 3339* (*p*_*discovery*_ = 9.99 × 10^−12^; *p*_*replication*_ =1.77× 10^−27^) and *Solobacterium uSGB 315* (an anaerobic gram-positive bacterium associated with colorectal cancer^25^; *p*_*discovery*_ = 2.12 × 10^−11^; *p*_*replication* =_1.44 × 10^−31^). *APPL2* encoded a multifunctional adapter protein that binds to various membrane receptors, nuclear factors and signaling proteins to regulate many processes, such as cell proliferation, immune response, endosomal trafficking and cell metabolism. *APPL2*-associated three taxa all positively correlated with high sugar/fat dietary frequency (**Fig. 2b**; Spearman *p*= 3.18× 10^−6^ for *Prevotella jejuni, p= 1*.*33*× 10^−5^ *for Oribacterium uSGB 3339* and *p*= 6.43× 10^−9^ for *Solobacterium uSGB 315*) when checking the correlation between oral taxa and phenotypic traits in this cohort, in line with the role of *APPL2* in controlling glucose-stimulated insulin secretion^26^. Appl2 protein had been reported to play a negative regulatory role in inflammation^27^. Its associated three taxa also correlated with decreasing risk of dental calculus and gingival bleeding (**Fig. 2c;** Spearman p<0.001), thereby supporting a link between genetic variation in the *APPL2* gene, immune response and the abundance of these taxa.

**Fig. 2.**
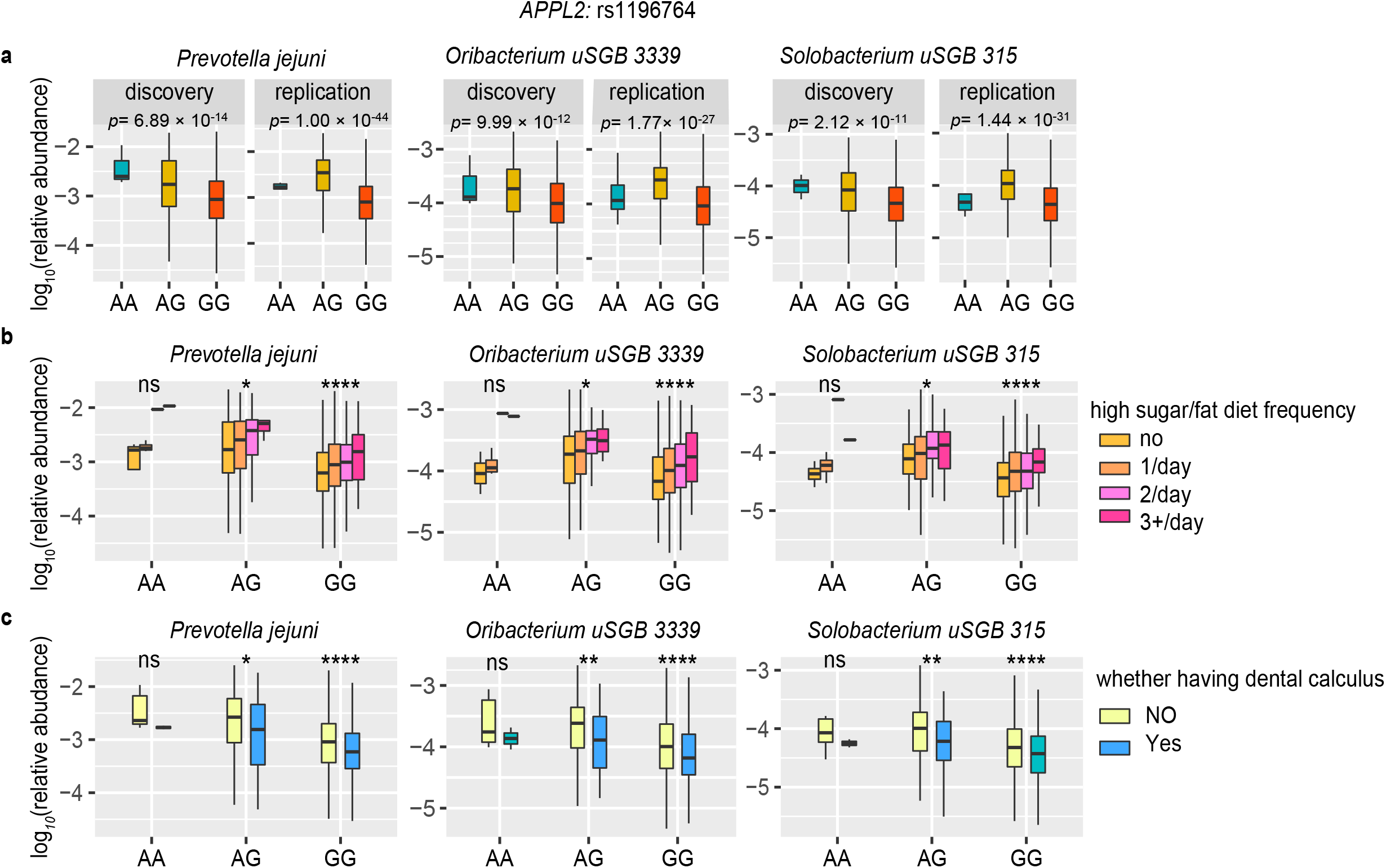
The links among human *APPL2* locus (rs1196764), three tongue dorsum bacteria, and diet as well as status of dental calculus. (**a**) The 3 panels present the associations of *APPL2* variation with microbial abundances of the three most significantly associated taxa: *Prevotella jejuni* (*p*_*discovery*_ = 6.89 × 10^−14^; *p*_*replication*_=1.00 × 10^−44^), *Oribacterium* uSGB 3339 (*p*_*discovery*_ = 9.99 × 10^−12^; *p*_*replication*_=1.77× 10^−27^) and *Solobacterium* uSGB 315 (*p*_*discovery*_ = 2.12 × 10^−11^; *p*_*replication*=_1.44 × 10^−31^); which reached study-wide significant in both discovery (N=2,017) and replication (N=1,333) cohorts. (**b**) Relative abundance of three taxa across stratified groups of individuals according to *APPL2*: rs1196764 variation and self-reported high sugar/fat dietary frequency (colored by no intake, 1/day; 2/day; 3+/day, respectively). (**c**) Relative abundance of three taxa across stratified groups of individuals according to *APPL2*: rs1196764 variation and whether having dental calculus (yellow: no and blue: yes). All statistical comparisons in (**b**) and (**c**) denote the p-values of Wilcoxon rank test on the total 3,350 individuals with log-transformed relative abundances. The significant code of p-values thresholds are showed: ns:p>0.05; *:p≤0.05; **:p≤0.01; ***:p≤0.001; ****:p≤0.0001.

The second strongest association was on rs3775944, which is a perfect proxy for the exonic variant rs10939650 (r^2^ = 0.99) in *SLC2A9*. Minor alleles of *SLC2A9* locus negatively correlated with *Oribacterium uSGB 1215* (**Fig. 3a**; *p*_*discovery*_ = 5.09 × 10^−13^; *p*_*replication*_ = 1.92 × 10^−5^), *Oribacterium uSGB 489* (**Fig. 3b**; *p*_*discovery*_ = 8.55 × 10^−11^; *p*_*replication*_ =1.62 × 10^−4^) and *L. umeaens*e (**Fig. 3c**; *p*_*discovery*_ = 4.69× 10^−9^; *p*_*replication*_ = 0.04). *SLC2A9* is a urate transporter and *SLC2A9* polymorphisms have been reported associated with serum uric acid and urine uric acid concentration in multiple studies^28-30^. We also looked at these top loci in Biobank Japan^31,32^, and *SLC2A9* was correlated with lower serum uric acid concentration (**Supplementary Fig. 4;** *p* = 5.56 × 10^−184^), ischemic stroke (*p* = 1.73 × 10^−4^), urolithiasis (*p* = 2.02 × 10^−4^) and pulse pressure (*p* = 6.86 × 10^−4^). The negative associations of *SLC2A9* with serum uric acid concentration (*p* = 6.74 × 10^−6^) and urine pH (*p* = 8.75 × 10^−4^) were confirmed in this cohort (**Supplementary Fig. 4)**. Interestingly, serum uric acid level highly correlated with *Oribacterium uSGB 1215* (**Fig. 3d**; Spearman rho = 0.27, *p* < 2.2×10^−16^).), *Oribacterium uSGB 489* (**Fig. 3e**; *rho = 0*.*24, p* < 2.2×10^−16^) and *L. umeaense* (**Fig. 3f**; *rho = 0*.*18, p* = 3.8 × 10^−10^). These results presented a potential explanation for *SLC2A9* acting on three oral taxa through serum uric acid as intermedium (**Fig. 3g**). Likewise, the lipoprotein lipase (*LPL) gene* was a determinant of triglyceride concentration (**Supplementary Fig. 4;** *p* = 5.93 × 10^−56^), triglyceride concentration correlated with abundance of *Haemophilus D parainfluenzae A* (*p* = 7.70 × 10^−16^), and consistently *LPL* exhibited significant association with *Haemophilus D parainfluenzae A* (*p* = 1.59 × 10^−8^). These findings suggested that host genes may regulate oral microbiota by mediating their relevant metabolites.

**Fig. 3.**
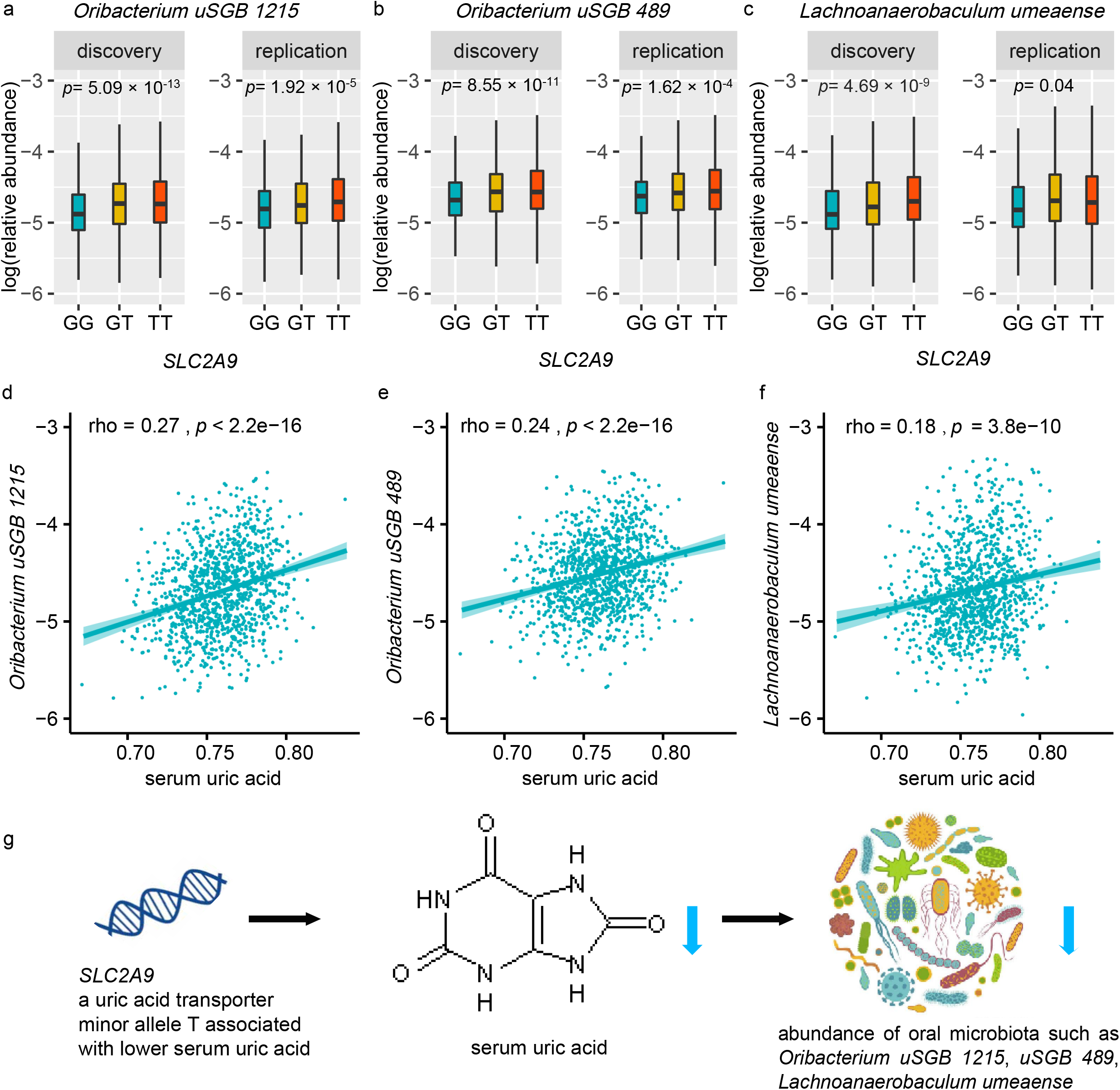
Interaction of human *SLC2A9* locus (rs3775944), serum uric acid, and three tongue dorsum bacteria. (**a**) The 3 panels present the associations of *SLC2A9* variation with microbial abundances of the three most significantly associated taxa: *Oribacterium* uSGB 1215 (**Fig. 2a**; *p*_*discovery*_ = 5.09 × 10^−13^; *p*_*replication*_= 1.92 × 10^−5^), *Oribacterium* uSGB 489 (**Fig. 2b**; *p*_*discovery*_ = 8.55 × 10^−11^; *p*_*replication*_=1.62 × 10^−4^) and L. umeaense (**Fig. 2c**; *p*_*discovery*_ = 4.69× 10^−9^; *p*_*replication*_= 0.04); which reached study-wide significant in discovery (N=2,017) cohort and well replicated in replication (N=1,333) cohort. (**b**) Correlations between three tongue dorsum bacteria and serum uric acid. Spearman’s correlation coefficient (rho) and p-value were showed. (**c**) Schematic representation of the interaction among *SLC2A9* locus (rs3775944), serum uric acid, and three tongue dorsum bacteria: *SLC2A9* as a uric acid transporter, its minor allele T of SNP rs3775944 associated with lower serum uric acid, and genetic predisposition to lower serum uric acid level is associated with lower abundance of *Oribacterium uSGB 1215, Oribacterium uSGB 489* and *Lachnoanaerobaculum umeaense*.

Variants in *MGST1* (leading SNP rs7294985) were identified as the third strongest signal, with minor allele negatively associated with *Streptococcus uSGB 2460* (*p*_*discovery*_ = 1.50 × 10^−11^; *p*_*replication*_ = 0.92), followed by family Streptococcaceae and other eight *Streptococcus* SGBs, such as *S. infantis* and *S. pseudopneumoniae*, although not confirmed in replication cohort. These variants were also positively associated with red blood cell count (*p* = 2.51 × 10^−5^) and asthma (*p* = 5.03 × 10^−5^) in Biobank Japan. Consistently, 84% (237/282) of the *Streptococcus spp*. were observed correlated with red blood cell count (*p* < 0.05), such as *S. mitis (p=* 1.99 × 10^−12^) and *S. pseudopneumoniae (p =* 4.51 × 10^−12^). These results suggested that commensal *Streptococcus* species might utilize red blood cells as camouflage to avoid being engulfed by phagocytic immune cells in addition to the well-known group A Streptococcus (*S. pyogenes*)^33^. Our results also supported previous findings that *Streptococcus spp*. are often involved in diseases of the respiratory tracts such as asthma^34^.

In addition to the above three study-wide significant loci, other well replicated genome-wide significant associations included rs17070896 in *ADAMTS9* with *Simonsiella muelleri*, rs59134851 near *MST1L-MIR3675* with *Streptococcus anginosus* (playing important roles in respiratory infections^35^), chr22:41198300 in *EP300-AS1* with *Parvimonas micra* (potential pathogen of colorectal cancer^36^), rs34555647 near *MIR3622B-CCDC25* with *Selenomonas sputigena* (potential pathogen of periodontal diseases^37^). These replicated associations invited further investigation of the impaction of host-microbial interactions on disease.

### M-GWAS of the salivary microbiome confirm and extend human genetic contribution to the oral microbiome

The saliva may appear more dynamic than the tongue dorsum, and the microbiome composition involves multiple niche in the oral cavity^38^. We next tried M-GWAS analysis for the saliva microbiome. With the 1,685 independent salivary microbial taxa (r^2^ < 0.8 from 3,677 taxa total), and 10 million human genetic variants (MAF ≥ 0.5%) in discovery cohort, 466 independent associations involving 374 independent loci (r^2^ < 0.2) reached genome-wide significance (*p* < 5 × 10^−8^). With a more conservative *Bonferroni*-corrected study-wide significant p-value of 2.97× 10^−11^ (= 5 × 10^−8^ / 1,685), 2 study-wise significant independent loci were identified (**Fig. 4a**). Similar to tongue dorsum M-GWAS analyses, the genomic inflation factors of these salivary M-GWAS tests showed no inflation (λ_GC_ ranged from 0.978 to 1.022 with median 1.002; **Supplementary Fig. 3b**). All genome-wide significant associations were listed in **Supplementary Table 3**.

**Fig. 4.**
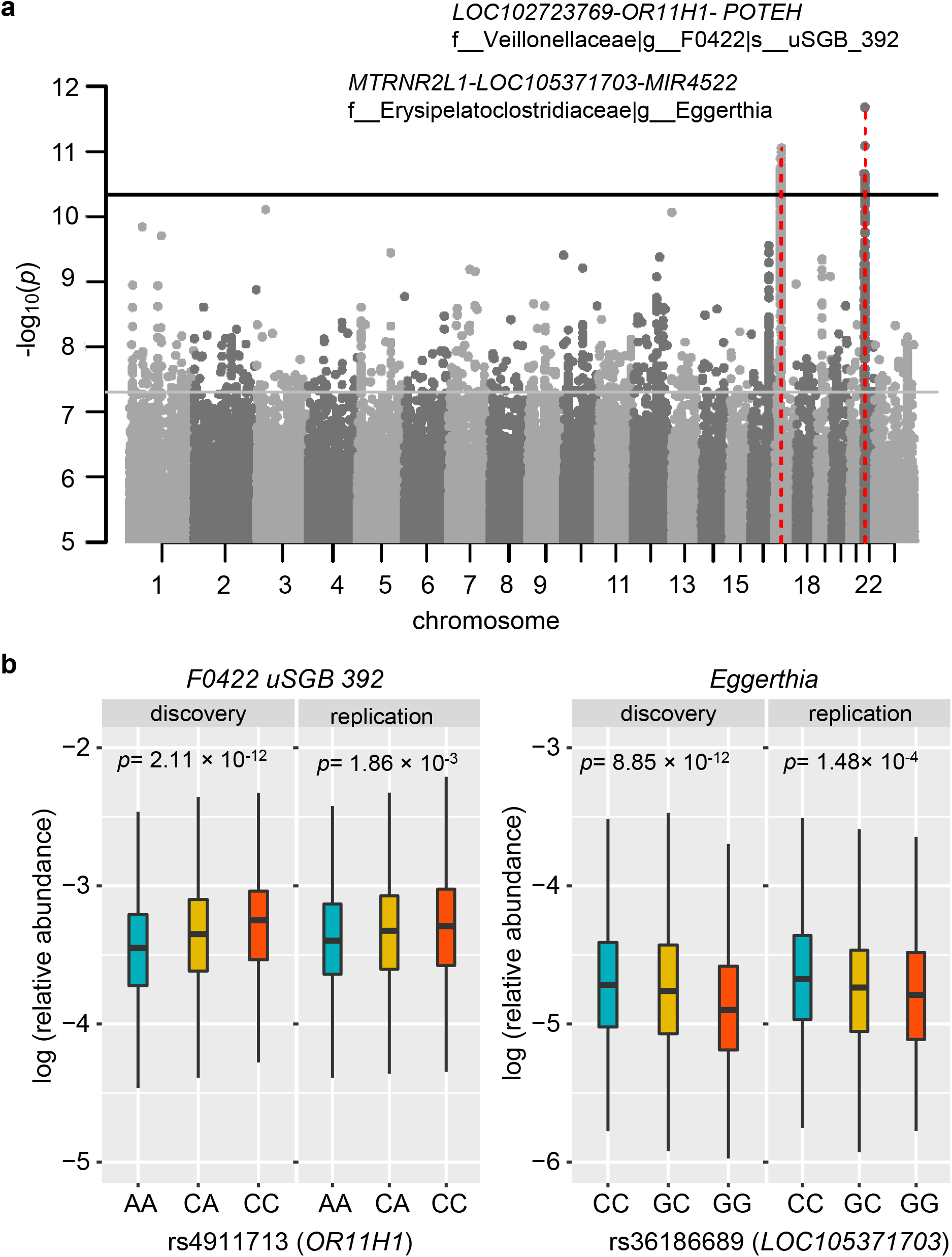
Host genetic signals associated with salivary microbiome. (**a**). Manhattan plot shows the genetic variants associated with the salivary microbial taxa. The horizontal grey and black lines represent the genome-wide (*p*□=□5□×□10^−8^) and study-wide (*p*□=□2.97□×□10^−11^ for 1,685 independent M-GWAS tests) significance levels, respectively. Two loci that associated with salivary microbiome and reached study-wide significance were marked in red. Their located genes and associated microbial taxa with *p* values of < 2.97□×□10^−11^ were also listed. (**b**)

As for validation, we were able to replicate 28 of the remaining 443 associations in the same effect direction of minor allele (*p* < 0.05), given that 23 of the 466 independent associations identified in the discovery cohort with *p* < 5 × 10^−8^ were not available in the low-depth replication dataset **(Supplementary Table 3)**. Two study-wide significant signals identified by this saliva M-GWAS were both well replicated (**Fig. 4b**). One genetic locus, spanning three genes *LOC102723769, OR11H1* and *POTEH*, associated with species *F0422 uSGB 392* belonging to family Veillonellaceae (leading SNP rs4911713; *p*_*discovery*_ = 2.11 × 10^−12^; *p*_*replication*_ =1.86 × 10^−3^). *F0422 uSGB 392* negatively correlated with concentrations of serum amino acids such as cystine and glycine and levels of blood microelements such as magnesium and lead, as well as serum testosterone levels and mental distress (impatience or tension) (**Supplementary Fig. 5**). The locus was consistently associated with mental distress (impatience) and serum testosterone level (p<0.05), while searching GWAS summary statistics from Biobank Japan and this study. The other locus, *MTRNR2L1-LOC105371703-MIR4522*, associated with genus *Eggerthia* (leading SNP rs36186689; *p*_*discovery*_ = 8.85 × 10^−12^; *p*_*replication*_ = 1.48× 10^−4^). *Eggerthia* was most positively associated with frequency of gingival bleeding, dental calculus, frequency of tooth pain and dental periodontitis, but negatively linked to serum hormones such as cortisone, aldosterone and testosterone (**Supplementary Fig. 6**). The locus was most associated with glaucoma and serum creatine level, while searching GWAS summary statistics from Biobank Japan and this study. Notably, the two loci both regulated the expression of genes in testis or brain cerebellar hemisphere (*p* <10^−5^) when searching in GTEx^39^ database, and their associated oral taxa both consistently correlated with serum testosterone levels and mental distress (impatience)(**Supplementary Figs. 5-6**). In addition, we found four loci associated with both salivary microbiome and metabolic traits or diseases at genome-wide significance: *DPEP2/NFATC3* that associated with species *Lancefieldella sp000564995* was linked to high density lipoprotein cholesterol (HDLC); *PDXDC2P-NPIPB14P* associated with species *Centipeda sp000468035* linked to thyroid abnormality; *LARP1* associated with species *Aggregatibacter kilianii* linked to mean corpuscular hemoglobin; *SMARCA1* associated with species *Veillonella parvula* linked to pharyngeal mucosal congestion (PMC) (**Supplementary Fig. 7**). These results were again confirmed a host genes - blood metabolites - oral microbiota axis in human body.

Among 455 and 466 independent associations identified for tongue dorsum and salivary microbiome (*p* < 5 × 10^−8^), respectively, 6 were shared between them (**Fig. 5**): *APPL2* associated with *Oribacterium uSGB 3339, LOC105374972-NRSN1* associated with *Lancefieldella uSGB 2019*; *CCL26-CCL24* associated with *Treponema B uSGB 706*; *RALGPS2* associated with *Scardovia wiggsiae*; *KRT16P1-LGALS9C* associated with *Patescibacteria uSGB 2650*; and *RTTN-SOCS6* associated with *Firmicutes uSGB 1705*. More specifically, among the 455 independent and genome-wide significant variants-taxa associations for the tongue dorsum samples, 386 associations (85%) were replicated with *p* < 0.05 in the same effect direction of minor allele for the salivary samples **(Fig. 5a-b)**. For example, *SLC2A9*, a determinant of low uric acid (UA) concentration, showed the strongest association with SGBs belonging to *Oribacterium* (*p* = 5.09 × 10^−13^) and *Lachnoanaerobaculum* (*p* = 4.69 × 10^−9^) in tongue dorsum samples, and also relative low association with that of *Oribacterium* (*p* = 0.001) and *Lachnoanaerobaculum* (*p* = 1.0 × 10^−4^) in salivary samples. Among the 466 independent and genome-wide significant variants-taxa associations for saliva samples, 391 associations (84%) also were replicated with p < 0.05 in the same effect direction of minor allele for the tongue dorsum **(Fig. 5c-d)**, although the top two study-wide significant loci for saliva didn’t reach suggestive significance in tongue dorsum samples (*p* > 1 × 10^−5^). Our M-GWAS of the salivary microbiome further confirm and extend human genetic contribution to the oral microbiome. These results suggested tongue and salivary microbiome as niches in one oral cavity shared high level of host genetic similarity in co-evolution process.

**Fig. 5.**
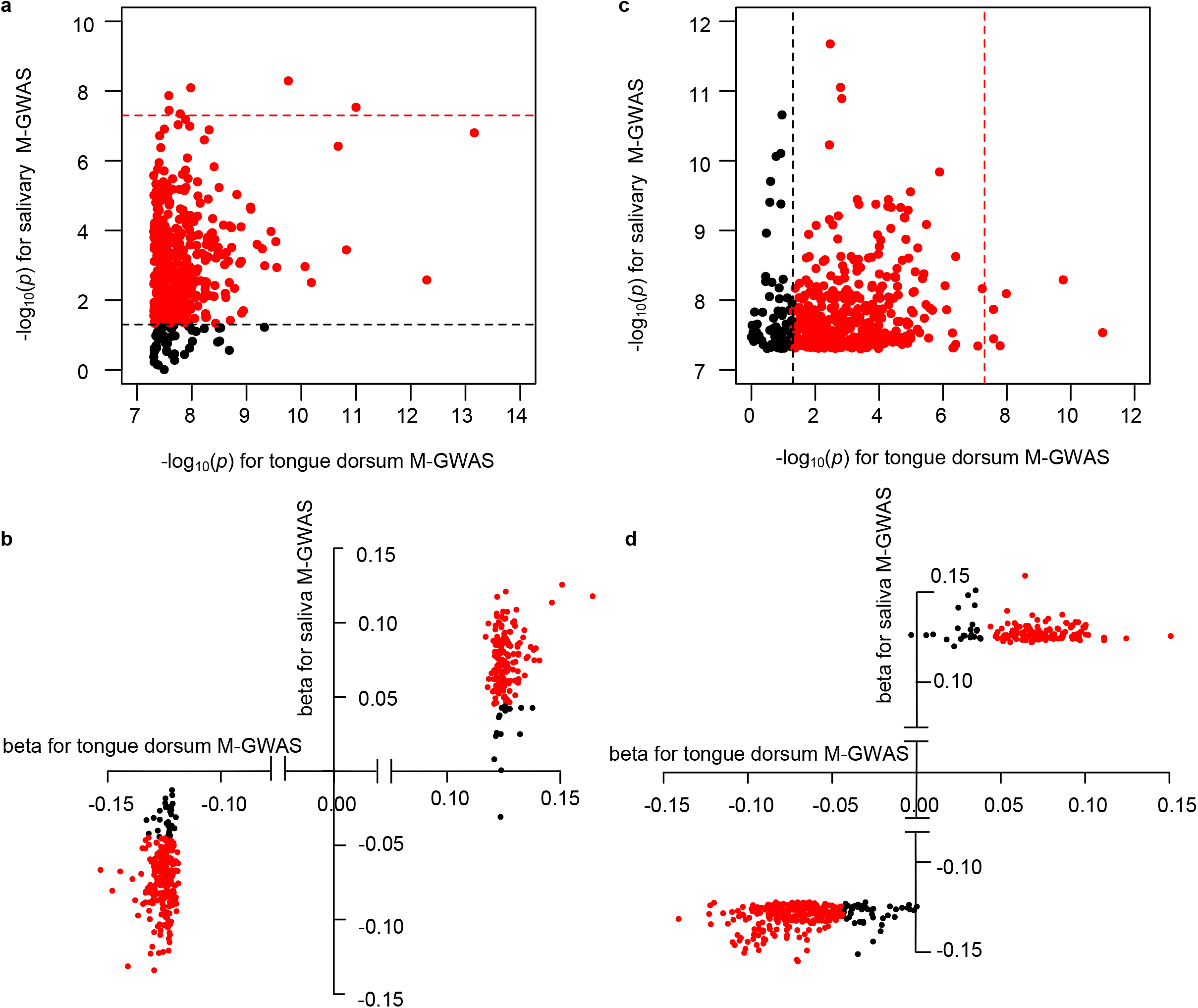
Comparisons of M-GWAS associations between tongue dorsum and saliva samples. (**a**). *p*-values comparisons of the 345 and 374 independent loci associated with tongue dorsum and salivary microbiome (p < 5 × 10^−8^), respectively. (**b**) β-values comparisons of the 345 and 374 independent loci associated with tongue dorsum and salivary microbiome (p < 5 × 10^−8^), respectively. In (a) and (b), the 6 genome-wide significant loci shared by tongue dorsum and salivary microbiome were listed. The dots marked red and black represented shared associations by both (P<5 × 10^−8^ in one niche and p<0.05 in the other niche in the same direction of minor allele) and specific associations (P<5 × 10^−8^ in one niche but p>0.05 in the other niche or in the opposite direction of minor allele) in one niche, respectively.

### Gene set enrichment analysis for oral M-GWAS signals

To explore the potential functions of the identified M-GWAS signals for tongue dorsum and salivary, we annotated the genetic associations and performed functional mapping and gene sets enrichment analysis with the DAVID^40^ and FUMA^41^ platform (Methods), followed by disease enrichment and tissue expression analysis. M-GWAS analysis returned 221 and 261 genes (<20kb for associated genetic loci) for tongue dorsum and salivary microbiome, respectively. Functional mapping of their separately related genes in DAVID database suggested that tongue dorsum associated host genes mainly enriched in phosphatidylinositol-related pathways including phosphatidylinositol signaling system, biosynthesis, dephosphorylation and phosphatidylinositol-3,4,5-trisphosphate 5-phosphatase activity, and Ca^2+^ pathway including calcium ion binding, calcium channel regulator activity and voltage-gated calcium channel activity (**Supplementary Table 4**). Phosphatidylinositol signaling system have been reported to be higher in the gut microbiota of centenarians^42^ and consistently decreased in saliva microbiota of RA patients^43^. Saliva associated host genes mainly enriched in cardiomyopathy including arrhythmogenic right ventricular-, hypertrophic- and dilated cardiomyopathy, glycerophospholipid metabolism and choline metabolism in cancer (**Supplementary Table 5**).

The GAD_Disease (Genetic Association Disease Database) segment analysis in DAVID showed that both tongue dorsum and saliva M-GWAS signals were enriched in cardiometabolic diseases and traits such as tobacco use disorder, myocardial infarction, triglycerides, blood pressure, lipoproteins, coronary artery disease, and nervous system diseases such as schizophrenia, bipolar disorder, psychiatric disorders and Parkinson’s disease (**Supplementary Tables 4 and 5**). Positional mapping in GWAS catalog using FUMA tool showed the similar diseases enriched results with that of using GAD catalog in DAVID. Genotype-Tissue Expression (GTEx) analysis on saliva microbiome associated host genes exhibited an enrichment for genes expressed in brain (anterior cingulate cortex BA24 and substantia nigra) and cells of EBV-transformed lymphocytes (**Supplementary Fig. 8**).

### Host genetics influence oral microbiome more than environment

We first investigate the contribution of environmental factors to oral microbiome β-diversity (based on genus-level Bray–Curtis dissimilarities), by using host metadata including age, gender, BMI, dietary, lifestyle, drug use and health status questions, as well as blood measurements. We selected 340 independent variables out of the total 423 environmental factors for association analysis (correlation r^2^ < 0.6). A total of 35 and 53 factors were significantly associated with β-diversity (BH-adjusted FDR□<□0.05) for tongue dorsum and salivary samples, respectively, via PERMANOVA analysis (**Supplementary Fig. 10; Supplementary Tables 6 and 7**). Of these, high sugar and high fat food frequency and dental calculus were the strongest associated factors for both tongue dorsum and salivary microbial compositions. A high sugar diet increased the abundances of some specific bacteria such as *Streptococcus mutans* that metabolized sugar to acids and caused dental caries. In this cohort, high sugar and high fat food frequency significantly increased the abundances of *Gemella haemolysans* (β = 0.21; *p* =2.92 × 10^−19^) and *Streptococcus parasanguinis* (β = 0.18; *p* = 7.56 × 10^−16^) in salivary samples. In total, 35 and 53 factors were able to infer 6.36% and 7.78% of the variance of microbiome β-diversity for tongue dorsum and salivary samples, respectively. When calculating the cumulative explained variance of β-diversity by using all the independent environmental variables, we found that 12.85% and 15.54% of the variance can be explained for tongue dorsum and salivary samples, respectively.

We next evaluated the effect of host genetics on oral microbiome compositions. We performed association analysis for α-diversity and β-diversity using 10 million genetic variants (MAF ≥ 0.5%). Six genome-wide significant loci were identified for α-diversity for oral microbiome (**Supplementary Table 8**). Four loci, *NFIB, LINC02578, LOC105373105* and *EIF3E*, associated with α-diversity of tongue dorsum samples. Two loci, *SLC25A42* and *LINC02225*, associated with α-diversity of salivary samples. In the association analysis between genetic variation and microbiome β-diversity, we found one locus for tongue dorsum samples and one locus for salivary samples with marginal genome-wide significance (*p* < 5 × 10^−8^; **Supplementary Fig. 11**), respectively. One SNP, rs545425011 located in *DNAJC12* was associated with microbial composition of tongue dorsum (*p* = 1.07 × 10^−8^). When searching its correlations with microbial taxa, it was mostly negatively associated with *Leptotrichia A sp000469505* and *Prevotella saccharolytica* (**Supplementary Table 9**), however, positively associated with *Rothia* SGBs such as *R. mucilaginosa which* was dominant in tongue dorsum and often observed in large patches toward the exterior of the consortium. The other SNP, rs73243848 located in *G2E3-AS1* was associated with salivary microbial composition (*p* = 2.35 × 10^−8^). It was mostly positively associated with *Prevotella* uSGB 2511 and family Bacteroidaceae (**Supplementary Table 10**).

The above analysis found that 53 and 35 factors (BH P <0.05) explained 7.78% and 6.36% of the β-diversity variance for salivary and tongue dorsum microbiome, respectively. By applying the same number of SNPs that were most closely associated with β-diversity, we identified 14.14% and 10.14% of the β-diversity variance could be inferred for salivary and tongue dorsum microbiome, respectively (**Supplementary Fig. 12**). The findings suggested host genetics is likely to influence oral microbiome more than environment.

### Host genetics and oral microbiome predict dental diseases

The dynamic and polymicrobial oral microbiome is a direct precursor of diseases such as dental calculus and gingival bleeding. To understand the aggregate effect of host genetic variants and oral microbiome on dental diseases, we constructed models using genetic polygenic risk scores (PRS) and oral microbiome separately, as well as their combination, to predict dental diseases. We found 2 of the 6 dental diseases occurred in over 5% individuals to be significantly associated with the oral microbiome (**Fig. 6a**; FDR *p*< 0.001). Either of salivary and tongue dorsum microbiome explained 20% of the variance for dental calculus. Salivary and tongue dorsum microbiome explained 13% and 15% of the variance for gingival bleeding, respectively. Compared with oral microbiome, the genetic PRS showed significantly higher predictive efficiency with a mean R^2^ of 45%, ranging from the lowest of 25% for gingival bleeding to highest of 60% for teeth loss. Furthermore, when incorporating the oral microbiome into PRS model, the predictive efficiency is slightly improved, with a 4% increasement of R^2^ for dental calculus and a 6% increasement of R^2^ for gingival bleeding (**Fig. 6b**).

**Fig. 6.**
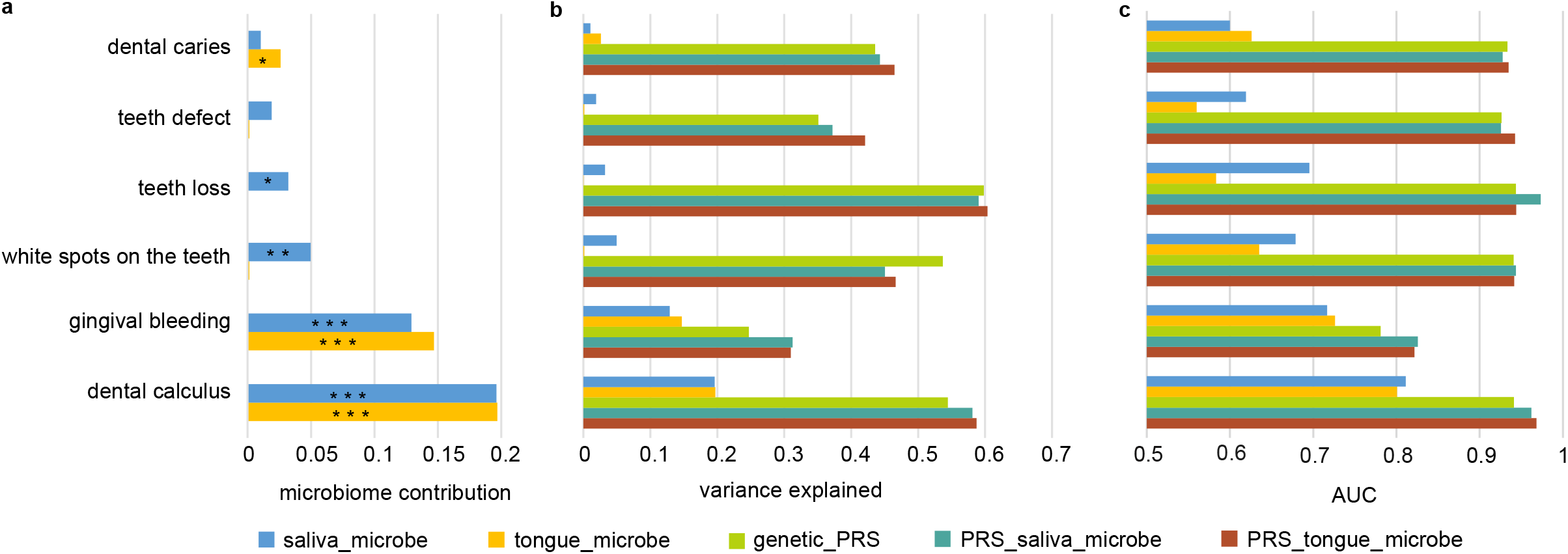
Oral microbiome and genetic PRS infer a significant fraction of the variance of dental diseases. (**a**) R^2^ estimates of six dental diseases and their significance contributed by oral microbiome, evaluated using linear model in lightGBM package. * *p* < 0.05, ** *p* < 0.01 and *** *p* < 0.001. (**b**) Predictive efficiency of six dental diseases (measured using coefficient of determination (R2)), evaluated using a linear model under five different sets of predictive features: (i) relative abundances of salivary microbial taxa; (ii) relative abundances of tongue dorsum microbial taxa; (iii) PRS calculated as an unweighted sum of risk alleles from independent and significant SNPs (LD r^2^ < 0.2, *p* < 10^−5^) for each oral disease; (iv) ‘PRS + salivary microbiome’: PRS, relative abundances of salivary microbial taxa and (v) ‘PRS + tongue microbiome’: PRS, relative abundances of tongue dorsum microbial taxa. (**c**) The discriminative efficiency for six dental diseases (measured using area under the curve (AUC)), evaluated using a discriminative model under five different sets of predictive features as described in (**b**).

The discriminative efficiency for dental diseases was also evaluated using area under the curve (AUC; **Fig. 6c**). Salivary and tongue dorsum microbiome had a good discrimination for dental calculus (AUC=0.81 and 0.80, respectively), and a median discrimination for gingival bleeding (AUC=0.72 and 0.73, respectively). The models of PRS had AUC of 0.93-0.94 for 5 of the 6 dental diseases, except for gingival bleeding (AUC=0.78). Incorporating the oral microbiome into PRS model resulted in improved discrimination with AUC increasing from 0.94 to 0.97 for dental calculus and from 0.78 to 0.83 for gingival bleeding. These results may help explain why some people are genetically predisposed to the major dental diseases.

## Discussion

In summary, we performed the first large-scale M-GWAS for oral microbiome and report unequivocal human genetic determinants for the oral microbiome. Based on the metagenome-assembled profile, we identified abundant genetic loci to associate with oral microbiota. 4 out of the 5 study-wise and nearly 1/10 genome-wide significant signals could be replicated in a low-depth genome cohort also from China, highlighting the power of adding independent cohort for genomic association analyses of oral microbiome. Our M-GWAS analysis found 84%-85% concordant association signals shared by tongue dorsum and salivary microbiome, with all genome-wide significant associations in one niche (**Fig. 5**; *p* < 5 × 10^−8^) were also at least nominally significant in the other niche (*p* < 0.05), consistent with our and previous findings that tongue dorsum and salivary microbiome communities exhibited high levels of similarity^44,45^, especially in micron-scale structure of oral niches^38,46^. Not only an independent cohort but also different niches (tongue dorsum and saliva) in the oral cavity corroborated the robust and replicable host-microbe association results. The non-replicated associations may require future more high-depth genome cohorts or other niches microbiome for further confirmation. Consistent with previous studies^24,47^, the salivary microbiome showed higher alpha diversity than tongue dorsum. In combination with the fact that saliva comes into contact with all surfaces in the oral cavity and represents a fingerprint of the general composition of the oral microbiome, these results suggested that salivary microbiome is more diverse and likely more dynamic. Thus, host genetic associations that are stronger with the salivary than the tongue dorsum community will further invite other omics studies, especially the proteome and the nitrogen cycle that could impact microbial growth.

Host-associated microbial communities are influenced by both host genetics and environmental factors. The debate centers on the relative contributions of host genetic and environmental factors to human microbiome. Twins modeling have demonstrated that some taxa of the human oral microbiome are heritable^18,19^, however, some studies indicated oral microbiome variances were shaped primarily by the environment rather than host genetics^20,21^. With this high-depth whole genome and metagenomic sequencing and high-quality assembled oral microbiome samples, we found that significant environmental factors explained 6.36%-7.78% of the β-diversity variance for oral microbiome, however, the same number of significant host SNPs could infer 10.14%-14.14% of the β-diversity variance for oral microbiome (**Supplementary Fig. 7 and 9**). These findings indicated host genetics is likely to influence oral microbiome more than environment. A previous study identified 42 SNPs that together explained 10% of the variance of gut microbiome β-diversity^16^. The comparable explained variances of host genetics on gut and oral microbiome supported the “oral–gut axis” that oral microbe transmission to and subsequent colonization of the large intestine is common and extensive among healthy individuals^3^.

As the genetics are already there at birth, oral hygiene would be more important for people who are more likely to develop dental diseases and beyond. Despite different aetiologias, dental calculus and gingival bleeding are both driven by a combined function of the oral microbiota and host factors. However, dental caries, teeth defect and loss were mainly determined by host genetics and less influenced by oral microbiome in this cohort. These results help us to better understand the pathogenic mechanisms and aided the design of personalized therapeutic approaches for different oral diseases. These results also provide a rational for repeatedly taking oral samples, to study the mostly stable human genome, long-term trends and short-term dynamics in the oral microbiome.

## Methods

### Study subjects

All the adult Chinese individuals in this cohort were recruited for a multi-omics study, with some volunteers having samples from as early as 2015, which would constitute the time dimension in ‘4D’. The cohort included 2,984 individuals with blood samples collected during a physical examination in 2017 in the city of Shenzhen and all these individuals were enlisted for high-depth whole genome sequencing (**Supplementary Table 1**). 3,932 (2,017 tongue dorsum and 1,915 saliva) oral samples from this cohort were newly collected for whole metagenomic sequencing between 2017 to 2018 (**Supplementary Table 1**). As for replication, blood samples were collected from 1,494 individuals, out of which 1,397 had tongue dorsum samples and 1,363 had salivary samples for metagenomic sequencing. The replication cohort was designed in the same manner but organized at smaller scales in multiple cities (Wuhan, Qingdao, etc.) in China. The protocols for blood and oral collection, as well as the whole genome and metagenomic sequencing were similar to our previous literature^5,23,48^. For blood sample, DNA was extracted using MagPure Buffy Coat DNA Midi KF Kit (no. D3537-02) according to the manufacturer’s protocol. Tongue dorsum and salivary samples were collected with MGIEasy kit. For salivary sample, a 2x concentration of stabilizing reagent kit was used and 2 mL saliva was collected. DNA of oral samples was extracted using MagPure Stool DNA KF Kit B (no. MD5115-02B). The DNA concentrations from blood and oral samples were estimated by Qubit (Invitrogen). 500 ng of input DNA from blood and oral samples were used for library preparation and then processed for paired-end 100bp sequencing using BGISEQ-500 platform^49^.

The study was approved by the Institutional Review Boards (IRB) at BGI-Shenzhen, and all participants provided written informed consent at enrolment.

### High-depth whole genome sequence for this cohort

2,984 individuals with blood samples were sequenced to a mean of 33x for whole genome. The reads were aligned to the latest reference human genome GRCh38/hg38 with BWA^50^ (version 0.7.15) with default parameters. The reads consisting of base quality <5 or containing adaptor sequences were filtered out. The alignments were indexed in the BAM format using Samtools^51^ (version 0.1.18) and PCR duplicates were marked for downstream filtering using Picardtools (version 1.62). The Genome Analysis Toolkit’s (GATK^52^, version 3.8) BaseRecalibrator created recalibration tables to screen known SNPs and INDELs in the BAM files from dbSNP (version 150). GATKlite (v2.2.15) was used for subsequent base quality recalibration and removal of read pairs with improperly aligned segments as determined by Stampy. GATK’s HaplotypeCaller were used for variant discovery. GVCFs containing SNVs and INDELs from GATK HaplotypeCaller were combined (CombineGVCFs), genotyped (GenotypeGVCFs), variant score recalibrated (VariantRecalibrator) and filtered (ApplyRecalibration). During the GATK VariantRecalibrator process, we took our variants as inputs and used four standard SNP sets to train the model: (1) HapMap3.3 SNPs; (2) dbSNP build 150 SNPs; (3) 1000 Genomes Project SNPs from Omni 2.5 chip; and (4) 1000G phase1 high confidence SNPs. The sensitivity threshold of 99.9% to SNPs and 98% to INDELs were applied for variant selection after optimizing for Transition to Transversion (TiTv) ratios using the GATK ApplyRecalibration command.

We applied a conservative inclusion threshold for variants: (i) mean depth >8×; (ii) Hardy-Weinberg equilibrium (HWE) *p* > 10^−5^; and (iii) genotype calling rate > 98%. We demanded samples to meet these criteria: (i) mean sequencing depth > 20×; (ii) variant calling rate > 98%; (iii) no population stratification by performing principal components analysis (PCA) analysis implemented in PLINK^53^ (version 1.9) and (iv) excluding related individuals by calculating pairwise identity by descent (IBD, Pi-hat threshold of 0.1875) in PLINK. No samples were removed in quality control filtering. After variant and sample quality control, 2,984 individuals (out of which 2,017 had matched tongue dorsum and 1,915 had matched salivary samples) with about 10 million common and low-frequency (MAF ≥ 0.5%) variants were left for M-GWAS analyses.

### Low-depth whole genome sequence for replicate cohort

1,494 individuals in replication cohort were sequenced to a mean of 9x for whole genome. We used BWA to align the whole genome reads to GRCh38/hg38 and used GATK to perform variants calling by applying the same pipelines as for the high-depth WGS data. After completing the joint calling process with CombineGVCFs and GenotypeGVCFs options, we obtained 43,402,368 raw variants. A more stringent process in the GATK VariantRecalibrator stage compared with the high-depth WGS was then used, the sensitivity threshold of 98.0% to both SNPs and INDELs was applied for variant selection after optimizing for Transition to Transversion (TiTv) ratios using the GATK ApplyRecalibration command. Further, we kept variants with less than 10% missing genotype frequency and minor allele count more than 5. All these high-quality variants were pre-phased using Eagle (v2.4.1)^54^ and then imputed using Minimac3 (v2.0.1)^55^ with our previous 1,992 high-depth WGS dataset^48^ as reference panel. We retained only variants with imputation info. > 0.7, Hardy-Weinberg equilibrium P > 10^−5^ and genotype calling rate > 90%. Similar to what we have done for discovery cohort, samples were demanded to have mean sequencing depth > 5×, variant call rate > 95%, no population stratification and no kinship. Finally, 1,430 individuals (out of which 1,333 had matched tongue dorsum and 1,299 had matched salivary samples) with 8.6 million common and low-frequency variants (MAF ≥ 0.5%) from replication cohort were left for association validation analysis.

### Oral metagenomic sequencing and quality control

Metagenomic sequencing was done on the BGISEQ-500 platform, with 100bp of paired-end reads for all samples and four libraries were constructed for each lane. We generated 19.18 ± 7.90 Gb (average ± standard deviation) and 19.90 ± 7.73 Gb raw bases per sample for tongue dorsum samples in discovery and replication cohorts, respectively (**Supplementary Table 1**). We also generated 13.64 ± 2.91 Gb and 13.66 ± 2.80 Gb raw bases per sample for salivary samples in discovery and replication cohorts, respectively. After using the quality control module of metapi pipeline followed by reads filtering and trimming with strict filtration standards (not less than mean quality phred score 20 and not shorter than 51bp read length) using fastp v0.19.463, host sequences contamination removing using Bowtie2 v2.3.564 (hg38 index) and seqtk65 v1.3, we finally got an average of 13.3Gb (host rate:31%) and 3.1Gb (host rate:77%) raw bases per sample for tongue dorsum and salivary samples, respectively.

### Oral metagenomic profiling

The high-quality oral genome catalogue was constructed in our previous study^23^. The oral metagenomic sequencing reads was mapped to oral genome catalogue (http://ftp.cngb.org/pub/SciRAID/Microbiome/human_oral_genomes/bowtie2_index) using Bowtie2 with parameters : “--end-to-end --very-sensitive --seed 0 --time -k 2 -- no-unal --no-discordant -X 1200”, and the normalized contigs depths were obtained by using jgi_summarize_bam_contig_depths, then based on the correspondence of contigs and genome, the normalized contig depth were converted to the relative abundance of each species for each samples. Finally, we merged all representative species relative abundance to generate a taxonomic profile for human oral population. The profiling workflow was implemented in metapi jgi_profiling module (https://github.com/ohmeta/metapi/blob/dev/metapi/rules/profiling.smk#L305).

### Tongue dorsum and salivary microbiome comparison

The nonparametric Wilcoxon rank-sum test was used to determine statistically significant differences in species α-diversity between tongue dorsum and saliva niches. We analyze the β-diversity (based on genus-level Bray–Curtis dissimilarity) difference between the two oral niches using PERMANOVA (adonis) in the ‘vegan’ package and visualize the two oral niches groups using ordination such as non-metric multidimensional scaling (NMDS) plots.

### Association analysis for oral microbial taxa

After investigating the distributions of occurrence rate and relative abundance of all microbial taxa, we decided to filter the microbial taxa to keep those with occurrence rate over 90% and average relative abundance over 1× 10^−5^. After filtering, the represented genera of these microbial taxa covered between 99.63% (tongue dorsum) and 99.76% (saliva) of the whole community in the cohort. As many oral microbial taxa are highly correlated and aims to reducing the numbers of GWAS tests, we then performed a number of Spearman’s correlation tests to obtain the independent taxa for M-GWAS analyses. Spearman’s correlations were calculated pairwise between all taxa, and the correlations used to generate an adjacency matrix where correlations of >0.8 represented an edge between taxa. A graphical representation of this matrix was then used for greedy selection of representative taxa. Nodes (microbiota taxa) were sorted by degree and the one with highest degree was then chosen as a final taxon (selecting at random in the case of a tie). The taxon and its connected nodes were then removed from the network and the process repeated until a final set of taxa sets were found such that each of the discarded taxon was correlated with at least one taxon. These filtering resulted in a final set of 1,583 and 1,685 independent microbial taxa for tongue dorsum and saliva, respectively, that were used for association analyses.

We tested the associations between host genetics and oral bacteria using linear model based on the relative abundance of oral bacteria. Specifically, the relative abundance was transformed by the natural logarithm and the outlier individual who was located away from the mean by more than four standard deviations was removed, so that the abundance of bacteria could be treated as a quantitative trait. Next, for 10 million common and low-frequency variants (MAF≥ 0.5%) identified in this cohort, we used a linear regression model to perform M-GWAS analysis via PLINK v1.9. Given the effects of environmental factors such as diet and lifestyles on microbial features, we included all potential cofounders that were significantly associated with the β-diversity (Benjamini–Hochberg FDR□<□0.05) estimates in the below explained variance analysis, as well as the top four principal components (PCs) as covariates for M-GWAS analysis in both the salivary and tongue dorsum niches.

To investigate the correlations between the identified oral microbiome-related SNPs and diseases, we downloaded the summary statistics data from the Japan Biobank^31,32^, a study of 300,000 Japanese citizens suffering from cancers, diabetes, rheumatoid arthritis and other common diseases. We searched the oral microbiome-related SNPs in the summary statistics data from Japan Biobank to examine their associations with diseases.

### Functional and pathway enrichment analysis

The significant genetic variants identified in the association analysis were mapped to genes using ANNOVAR^56^. Given that some significant genetic variants were low-frequency in the M-GWAS results, it’s most suitable to input gene lists for enrichment analysis. We mapped variants to genes based on physical distance within a 20kb window and got the gene lists for enrichment analysis. DAVID (https://david.ncifcrf.gov/) was utilized to perform functional and pathway enrichment analysis. DAVID is a systematic and integrative functional annotation tool for the analysis of the relevant biological annotation of gene lists and provide functional interpretation of the GO enrichment and KEGG pathway analysis^40^. The *p*-value <0.05 was considered statistically significant. In addition, the mapped genes were further investigated using the GENE2FUNC procedure in FUMA^41^ (http://fuma.ctglab.nl/), which provides hypergeometric tests for the list of enriched mapping genes in 53 GTEx tissue-specific gene expression sets, 7,246 MSigDB gene sets, and 2,195 GWAS catalog gene sets^41^. Using the GENE2FUNC procedure, we examined whether the mapped genes were enriched in specific diseases or traits in the GWAS catalog as well as whether showed tissue-specific expression. Significant results were selected if a false discovery rate (FDR)-corrected *p* < 0.05 was observed.

### Association analysis for microbiome α-diversity and β-diversity

The microbiome β-diversity (between-sample diversity) based on genus-level abundance data were generated using the ‘vegdist’ function (Bray–Curtis dissimilarities). Then, we performed principal coordinates analysis (PCoA) based on the calculated beta-diversity dissimilarities using the ‘capscale’ function in ‘vegan’. Finally, associations for β-diversity (a two-axis MDS) were performed using the manova() function from the ‘stats’ package, in a multivariate analysis using genotypes and the same covariates stated above as variables.

### Association analysis for environmental factors

As part of the 4D-SZ cohort, all participants in this study had records of multi-omics data, including anthropometric measurement, stool form, defecation frequency, diet, lifestyle, blood parameters, hormone, etc.^21^. A total of 423 environmental factors are available in this cohort. Environmental metadata were first log-transformed and checked for collinearity using the Spearman correlation coefficient. Collinearity was assumed if a Spearman’s ρ >□0.6 or ρ< −0.6. Collinear variables were considered redundant and one variable from each pair was removed from further analysis, resulting in a final set of 340 variables.

To investigate the potential associations of top loci identified in microbiome GWAS with environmental variables especially for serum metabolites, we also performed GWAS analysis for the 340 environmental variables. Among the 340 environmental traits, the log10-transformed of the mean-normalized values was calculated for each quantitative phenotype (such as amino acids, vitamins, microelements etc.) and a linear regression model for quantitative trait implemented in PLINK v1.9 was used for association analysis. Samples with missing values and values beyond 4 s.d. from the mean were excluded from association analysis. For each binary phenotype (such as diet, lifestyle etc.), a logistic regression model was used for association analysis. Age, gender and the top four PCs were included as covariates for each association analysis.

### Environmental factors explained variance of oral microbiome

We next searched for associations between the 340 environmental variables selected above and the oral microbiome compositions. We performed Bray-Curtis distance-based redundancy analysis (dbRDA) to identify variables that are significantly associated with β-diversity and measure the fraction of variance explained by the factors, using the ‘capscale’ function in the vegan package. The significance of each response variable was confirmed with an analysis of variance (ANOVA) for the dbRDA (anova.cca() function in the vegan package). Only the variables that were significantly associated (Benjamini–Hochberg FDR□<□0.05) with the β-diversity estimates in the univariable models were included in the multivariable model. The additive explanatory value (in %) of significant response variables (e.g. environmental parameters, vitamins and serum amino acids etc.) was assessed with a variation partitioning analysis of the vegan package (‘adj.r.squared’ value using RsquareAdj option).

### Construct PRS for diseases prediction

To obtain the predictions of human genetics on dental diseases, we used gradient boosting decision trees from the LightGBM (v.3.1.1) package^57^ implemented in Python (v3.7.8) and fivefold cross validation scheme to construct risk-prediction models. In every fold of the fivefold cross validation scheme, we calculated the associations between SNPs and dental diseases within the training dataset, and then selected independent and significant SNPs (LD r^2^ < 0.2, *p* < 10^−5^) to calculate the PRS as an unweighted sum of risk alleles, and finally we trained a model on the PRS and predicted the disease risk in test dataset. During the process, we obtained the optimal values of the tuning parameters using fivefold cross validation and evaluated the results using the coefficient of determination (R^2^) as variance explained and AUC as disease discriminative efficiency.

## Data availability

All summary statistics that support the findings of this study including the associations between host genetics and tongue dorsum microbiome, host genetics and saliva microbiome are publicly available from https://db.cngb.org/search/project/CNP0001664. The release of these summary statistics data was approved by the Ministry of Science and Technology of China (Project ID: 2021BAT1539). According to the Human Genetic Resources Administration of China regulation and the institutional review board of BGI-Shenzhen related to protecting individual privacy, sequencing data are controlled-access and are available via application on request (https://db.cngb.org/search/project/CNP0001664).

## Acknowledgments

We sincerely thank the support provided by China National GeneBank. We thank all the volunteers for their time and for self-collecting the oral samples using our kit.

## Author contributions

H.J. and T.Z. conceived and organized this study. J.W. initiated the overall health project. X.X., H.Y., S.Z., Y.H., Y.Zong and W.Liu contributed to organization of the cohort the sample collection and questionnaire collection. H.Lu led the DNA extraction and sequencing. X.Q., J.Z., R.W. generated the metabolic data. X.Liu, T.Z., and X.T. processed the whole genome data. Y.Zou, X.Lin, H.Liang, W.Li, Y.J., Y.Q. and L.Z. processed the metagenome data. X.Liu performed the metagenome-genome-wide association analyses. X.Liu and H.J. wrote the manuscript. All authors contributed to data and texts in this manuscript.

